# Quantifying bacterial fitness in intracellular dynamics^*^

**DOI:** 10.1101/539163

**Authors:** Francisco F. S. Paupério, Erida Gjini

## Abstract

Understanding bacterial infection is challenging because it involves a complex interplay of host, pathogen, and intervention factors. To design successful control measures, mathematical models that quantify such interplay at the level of populations and phenotypes are needed. Here, we study a key aspect of intracellular infection: the interaction dynamics between bacteria and target cells, applicable to pathogens such as *Salmonella*, *E. coli* or *Listeria monocytogenes*. Our mathematical model focuses on the macrophage-bacteria system, implicitly accounting for host immunity, and illustrates three infection scenarios driven by the balance between bacterial growth and death processes. Our analysis reveals critical parameter combinations for the intracellular vs. extracellular fitness advantage of persistent bacteria, and the drivers of overall infection success across acute and persistent regimes. Our results provide quantitative insights on transitions from persistent, to acute, to containment of infection, and suggest biological parameters, such as infected macrophage apoptosis rate and burst size, as suitable intervention targets.

## I. INTRODUCTION

Bacterial infections constitute a major concern for human health, especially in the context of rising antibiotic resistance [1], [2]. Clinical manifestation of infections caused by pathogenic bacteria reflects a complex interplay between microbe, host and intervention factors. While the innate and adaptive immune system actively fight infection, the growth dynamics of bacteria as a function of their resource is also very important. Macrophages are among the main phagocytic cell types of the innate immune system, and are often targeted by pathogenic bacteria for intracellular growth [3].

Several species have evolved sophisticated mechanisms to interfere with host factors for invasion, replication and spread in the intracellular compartment [4]. The intracellular lifestyle protects bacteria from the complement or adaptive immune response of the host, and from the activity of antibiotics [5]. Intracellular bacteria also compete less with other resident bacteria for nutrients. Bacterial species able to live within professional phagocytic cells such as macrophages include *Salmonella* [6], *Listeria monocytogenes* [7], *Mycobacterium tuberculosis* [8], *Shigella* and some *E. coli* strains. These can cause acute infection in their host or establish a persistent infection, sometimes in the absence of clinical disease.

In this study, we adopt a minimal mechanistic approach to describing and understanding intracellular infection dynamics of bacteria. What are the processes that determine acute versus persistent infection? How do the phenotypes at the host-pathogen interface interact? Here, we use a mathematical model to address these questions, and to capture the key features of infection dynamics driven by macrophage-bacteria interaction. Our model reveals critical combinations of bacterial life-history traits for fitness during infection and highlights the trade-off between intracellular growth and extracellular persistence. Our results should be relevant when comparing bacterial strains competing within host, variation in virulence, mutations conferring advantage in the intracellular vs. extracellular environment [9], antibiotics, and host-directed therapies [10].

## II. MATHEMATICAL MODEL

We study intracellular infection by modeling three populations: an extracellular bacterial population, *B*, target host cells representing uninfected macrophages, *M*, and infected macrophages, *I*, permissive to intracellular bacterial replication. The model shares features with previous models of bacterial infection [11], [12], [13] and viral dynamics models [14], [15], [16].

Extracellular bacteria infect susceptible macrophages at rate *β*, and then reproduce exclusively in the intracellular compartment, in macrophages. Upon necrosis at rate *δ*, infected phagocytes burst and release bacteria, at burst size *N*, in the extracellular environment. Infected macrophages can undergo apoptosis at rate *a*, in response to pro-inflammatory stimuli generated by the infection. Extracellular bacteria instead are eliminated in the host at rate, *c*, under the action of other immune cells from the innate immune system, e.g. neutrophils.

The role of adaptive immune mechanisms [6], [17] is assumed static, and can be considered implicit in this model, reflected possibly in the magnitude of parameters such as *β, a, c*. We assume for the baseline dynamics of uninfected macrophages a logistic growth with parameters *r*, and *K*, but the results are very similar if we were assume constant recruitment and decay rate under conservation of the same equilibrium level [11]. The dynamics are given by:

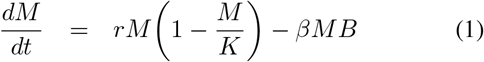

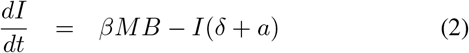

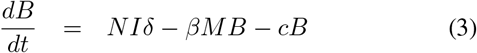

Notice, we do not model antibiotic treatment, as our aim is to provide analytic insights into the baseline dynamics at the local site of infection under no intervention. Initial conditions for the system are (*M*_0_, *I*_0_, *B*_0_) = (*K*, 100, 0). In addition, we assume an extinction threshold (*B*_*ext*_ = 0.1) to represent stochastic extinction at low bacterial population numbers. Model parameters and some values for illustration are summarized in Table 1.

**TABLE I.** MODEL PARAMETERS AND INTERPRETATION

| Parameter | Interpretation | Default values (i) | Units | Alternative values (ii) | Possible range | Reference |
| --- | --- | --- | --- | --- | --- | --- |
| $r$ | Growth rate of uninfected macrophages | 0.09 | $\text{cell}^{-1} \text{ day}^{-1}$ | 0.09 | $[0.01, 1]$ | [13] |
| $K$ | Maximum number of macrophages | $10^8$ | cells | $10^8$ | $[10^5, 10^8]$ | [11], [13] |
| $\beta$ | Infection rate of macrophages | $1.2 \times 10^{-7}$ | $\text{cell}^{-1} \text{ day}^{-1}$ | $1.2 \times 10^{-6}$ | $[10^{-8}, 10^{-6}]$ | [11], [13] |
| $\delta$ | Necrosis rate of infected macrophages | 0.2 | $\text{cell}^{-1} \text{ day}^{-1}$ | 0.9 | $[0.01, 20]$ | [11] |
| $a$ | Apoptosis rate of infected macrophages | 12 | $\text{cell}^{-1} \text{ day}^{-1}$ | 0.2 | $[0.01, 20]$ | [11], [12] |
| $N$ | Burst size of infected macrophages | 100 | cells | 4 | $[1, 100]$ | [6] |
| $c$ | Death rate of extracellular bacteria | 2 | $\text{cell}^{-1} \text{ day}^{-1}$ | 43 | $[0.1, 50]$ | [11] |

Although these do not represent a particular infection, they should fall within a realistic range that can be investigated further. We highlight their critical relationships, which hold in generic manner and are analyzed below.

## III. RESULTS

### A. Three infection outcomes

We find the model admits three possible infection outcomes: i) containment, when bacteria do not grow in the host but are gradually cleared, ii) growth, followed by persistent infection, iii) growth, followed by clearance yielding acute infection (Figure 1). The first scenario results when microbial death processes dominate over growth processes during infection, and replication inside macrophages is insufficient compared to clearance. For example, the extreme case *N* = 0 corresponds to total blocking of intracellular replication, and effective bacterial clearance through phagocytosis.

**Fig. 1.**
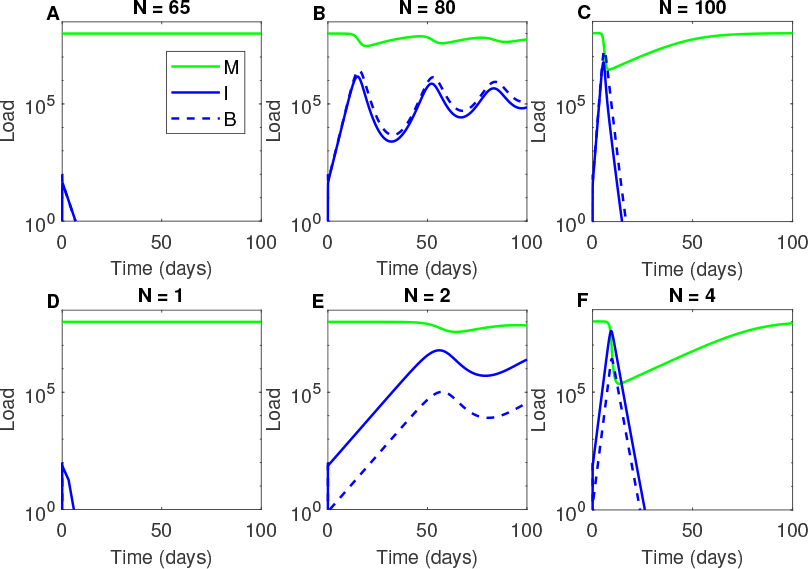
Three general scenarios of bacteria-macrophage dynamics. **A. D.** Containment of bacteria, clearance of infection. **B. E.** Growth and persistent infection. **C. F** Growth and acute infection followed by clearance (bacteria hit the extinction threshold). In this illustration, the burst size from infected macrophages, *N*, was varied to describe how bacterial replication capacity in host cells can produce big qualitative changes in the dynamics. Top row (A, B, C) other parameters as in Table 1(i). Bottom row (D, E, F) other parameters as in Table 1(ii).

The second scenario results when growth and death within host balance to maintain infection at intermediate levels. In that case, the macrophage-bacteria system undergoes a finely-tuned prey-predator type dynamics, characterized by oscillations around an equilibrium value. In this case, all three within-host populations remain in the positive range.

The third scenario, of an acute type, results when growth processes are very strong, leading to fast resource consumption followed by a drastic bacterial decline, such that extinction is hit before target cells have recovered. An extreme case of the third scenario results when both bacteria and its resource, in this case macrophages, go extinct due to bacterial over-proliferation.

We find that bacteria can persist within host only if the parameters satisfy a critical condition:

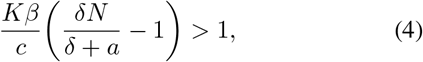

This is similar to the critical threshold for disease persistence given by the basic reproduction number, *R*_0_ > 1, in epidemic models [18]. Above, our condition says that only if the maximal level of resource, in this case the carrying capacity *K* of target host cells, is large enough relative to the ‘consumption’ rate, reflected in *β*, and such that growth dominates over death processes, the bacteria can persist. The reverse implies that the persistence equilibrium does not exist and that the containment equilibrium (*K*, 0, 0), with macrophages at their maximum level and no infection, is stable.

The exact expressions for the persistence equilibrium are given in the Appendix. Notice that an obvious sub-condition in Eq. 4 is a high enough intracellular replication of bacteria relative to the lifespan of infected macrophages: 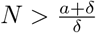.

In Figure 2 we illustrate how the persistence equilibrium depends on critical model parameters. Figure 2A shows that for a given apoptosis rate, the bacteria can persist only if burst size *N* is sufficiently big. The equilibrium level of extracellular bacteria decreases with burst size and apoptosis rate. Figure 2B shows how the bacterial peak depends on burst size, for three values of macrophage apoptosis rate. In Figure 2C-D, we show the dependence of theoretical equilibrium and numerical peak level of infected macrophages on model parameters.

**Fig. 2.**
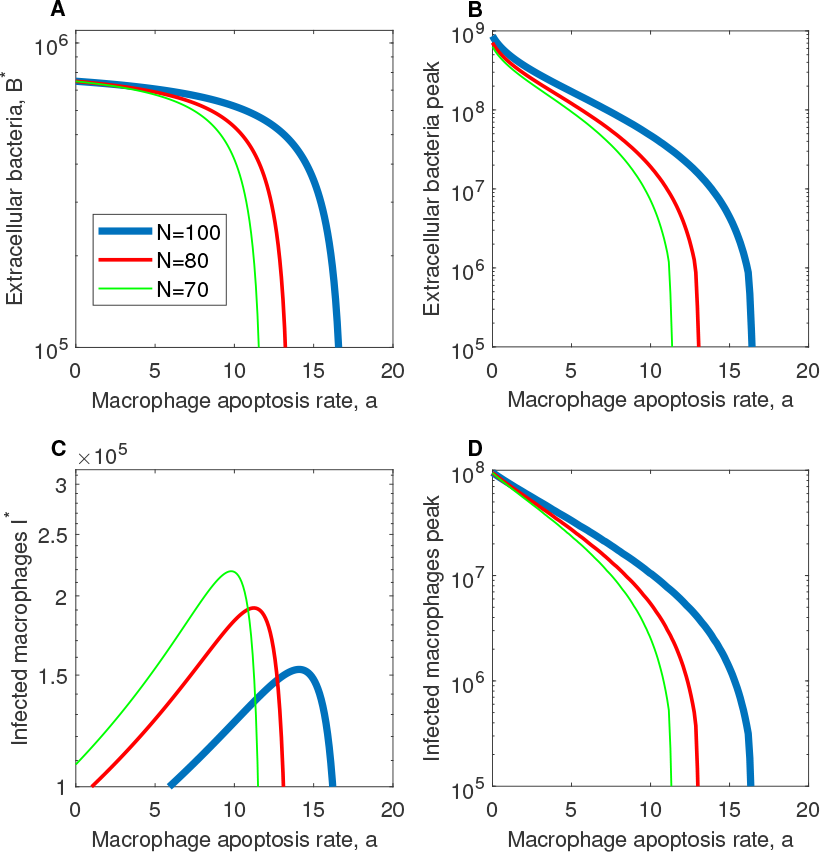
Bacterial persistence at theoretical equilibrium and peak infection severity as a function of model parameters. **A.** Analytical equilibrium level of extracellular bacteria *B^*^* is plotted as a function of *a*, for three different burst size values *N*. **B.** Peak bacterial load from simulations is plotted as a function of *a*: this corresponds to the maximum of *B*(*t*) reached in the first growth peak. **C.**Analytical equilibrium level of infected macrophages. **D.** Peak level of infected macrophages obtained numerically from simulations. Other parameters as in Table 1 (i).

As infected host cells are cleared faster, the opportunities for bacterial persistence decrease, leading to a lower and lower level of infection. This means in certain scenarios (e.g. *a* high), bacterial persistence can be maintained at undetectable levels, consistent with asymptomatic carriage.

### B. Intracellular vs. extracellular persistence

Our analytical approach to infection allows us to investigate more in detail the conditions for when intracellular microbial growth dominates bacteria found in the extra-cellular compartment. One important measure is the ratio between infected macrophages and extracellular bacteria at equilibrium, which in our model is given by:

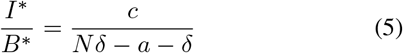

Figure 3 illustrates this ratio decreasing with *N*. extracellular bacteria start to dominate for high burst size values, and this happens at lower *N* when the lifespan of infected macrophages increases, i.e. for lower apoptosis rate *a*.

**Fig. 3.**
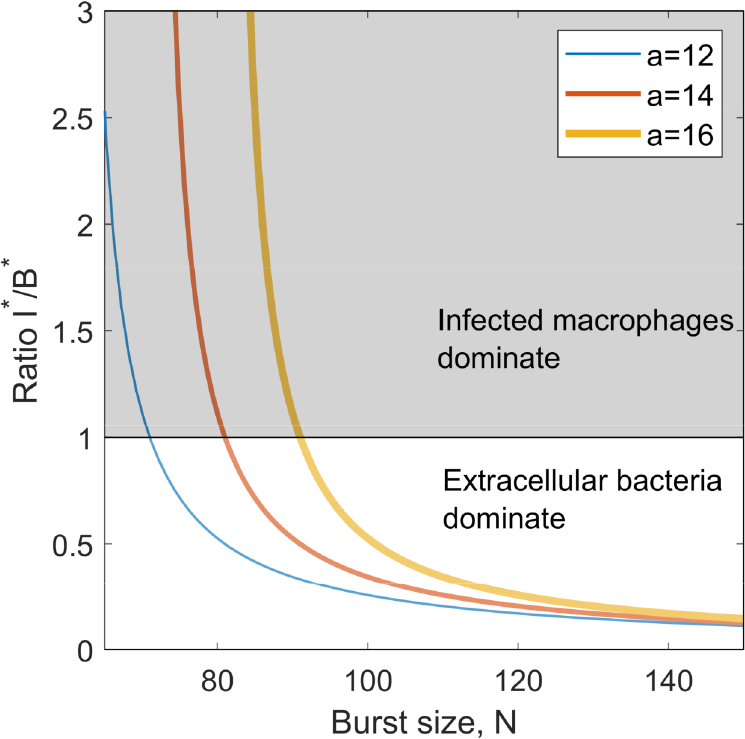
Intracellular vs. extracellular persistence. The ratio of infected macrophages and extracellular bacteria at persistence equilibrium is plotted as a function of burst size *N*. Other parameters as in Table 1 (i).

We must note however that the ratio *I^*^/B^*^* is more conservative with regards to extracellular bacteria, as it implicitly accounts for recently-infected macrophages, which start with one engulfed bacterium are more likely to harbour low cell numbers, compared to longer-lived (older) infected macrophages, which most likely harbour bacterial cell numbers close to *N*. But, since we do not track the age of infected cells in our model, we consider *I^*^/B^*^* to be a good indicator of intra-vs. extracellular persistence of bacteria.

Another measure for intracellular infection fitness is the proportion of macrophages that are infected at equilibrium: *I^*^/*(*I^*^* + *M ^*^*). Its explicit exact formula, in terms of model parameters, is more complicated, but in Figure 4 we illustrate for example how this quantity increases with burst size *N*, and *r*, the replenishment rate of uninfected macrophages. When *r* is 100-fold higher, the bacteria can maintain a roughly 100-fold higher proportion of infected cells. While fast replenishment of phagocytes may act as a strong positive feedback on infection, as apoptosis rate *a* increases, the proportion of infected macrophages goes down. Different parameters can have contrasting effects on infection.

**Fig. 4.**
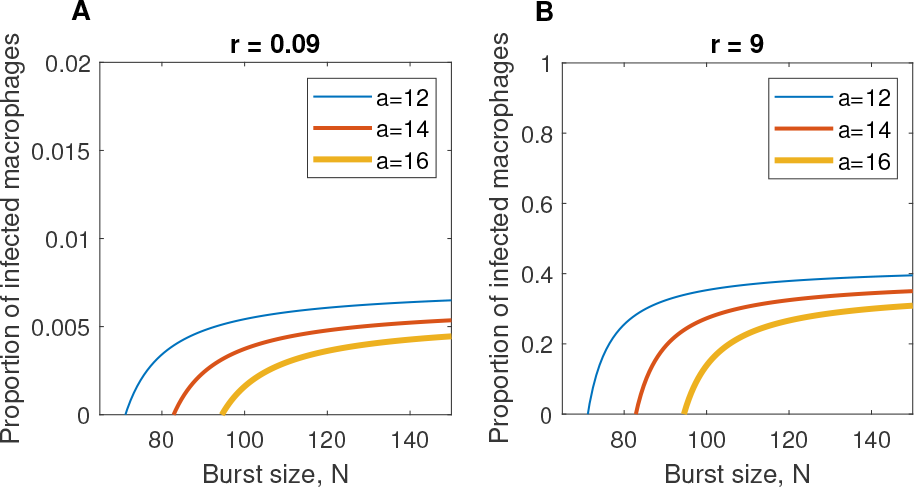
Proportion of infected macrophages in persistent infection. Equilibrium fraction of infected macrophages *I^*^/*(*I^*^* + *M^*^*) expected in persistent scenarios is plotted as a function of burst size for three values of infected macrophage apoptosis rate. **A.** low replenishment of uninfected target cells, **B.** high replenishment of target cells. Parameters in Table 1(i).

By taking advantage of our system’s analytical tractability, we also find an implicit expression for the proportion of infected macrophages, as a function of extracellular bacteria, given by a simple saturating function:

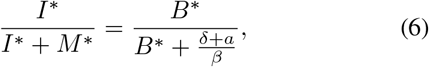

indicating that after a while, increasing the number of bacteria released in the extracellular environment (e.g. by higher *N*), does not pay off in terms of augmenting the prevalence of infection in macrophages.

## IV. DISCUSSION

In this study, we highlighted the main processes responsible for the outcomes of intracellular bacteria-macrophage dynamics. This system is regulated by parameters that control bacterial entry into target cells, proliferation and extracellular survival. For bacterial growth, a critical relationship between multiple fitness dimensions is needed, balancing intracellular growth and extracellular survival. Furthermore, only specific parameter combinations create optimal conditions for persistence. Too much proliferation may deplete the very conditions needed for growth, while not enough proliferation will make bacteria die out. Thus, an intermediate rate of growth seems optimal, as has been shown in studies which account for host mortality and pathogen virulence [19].

It is known that intracellular bacteria often display small-colony variants to increase within-host success [20], a trait shown to evolve during adaptation to intracellular lifestyle in macrophages [21]. Probably this reflects an optimal strategy to resolve the trade-off between ‘resource’ use for current proliferation and ‘resource’ availability for future growth, represented in our model, for example, with intermediate burst size. It is precisely at such attenuated growth rates that bacterial immune evasion is most probable, as the infection may persist undetected, hampering clearance.

Our model indicates that if any of the biological parameters changes during the course of time, an infected patient may shift from low-level chronicity to full-blown infection. The exact outcome will depend on the complex co-adaptation feedbacks between bacteria and host cells [22], and ultimately their net effect on cell population phenotypes. We provide several analytical insights into within-host constraints between multiple variables. These relationships can be used to integrate theory with empirical observations, such as those in [6], where one could try to estimate biological parameters from reported experimental variables.

Although only implicit in our model’s parameters, sufficient host immune competence is typically needed to reduce infection levels and drive bacteria towards ultimate and stable clearance. How microbial life-history traits respond to the presence of dynamic host defences, and eventually drug treatment feedbacks remains open to further investigation.

## ACKNOWLEDGEMENT

The work was supported by a NOS Alive-IGC fellowship to Francisco Paupério in 2017.

## V. APPENDIX

The system admits 3 equilibria: i) the trivial (0, 0, 0), the clearance (*K*, 0, 0), and the infection persistence (*M^*^, I^*^, B^*^*). The persistence equilibrium only exists if inequality (4) is satisfied and is given by:

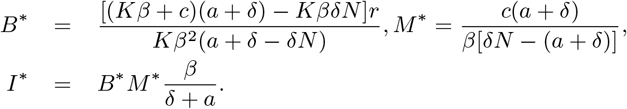

Thus, uninfected *M ^*^* are a linear function of extracellular bacteria: 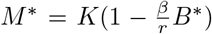, while infected macrophages *I^*^* are a quadratic function: 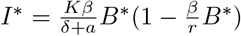.

## REFERENCES

[1] IM Gould. Antibiotic resistance: the perfect storm. International journal of antimicrobial agents, 34:S2–S5, 2009.

[2] WHO. Antimicrobial resistance: global report on surveillance 2014. 2014.

[3] Philippe Sansonetti. Phagocytosis of bacterial pathogens: implications in the host response. In Seminars in immunology, volume 13, pages 381–390. Elsevier, 2001.

[4] Chantal De Chastellier. The many niches and strategies used by pathogenic mycobacteria for survival within host macrophages. Immunobiology, 214(7):526–542, 2009.

[5] Françoise Van Bambeke and Maritza Barcia-Macay et al. Cellular pharmacodynamics and pharmacokinetics of antibiotics: current views and perspectives. Current Opinion in Drug Discovery and Development, 9(2):218, 2006.

[6] Denise M Monack, Donna M Bouley, and Stanley Falkow. Salmonella typhimurium persists within macrophages in the mesenteric lymph nodes of chronically infected nramp1+/+ mice and can be reactivated by ifnγ neutralization. Journal of Experimental Medicine, 199(2):231–241, 2004.

[7] José A Vázquez-Boland and Kuhn et al. Listeria pathogenesis and molecular virulence determinants. Clinical microbiology reviews, 14(3):584–640, 2001.

[8] Jean Pieters and John Gatfield. Hijacking the host: survival of pathogenic mycobacteria inside macrophages. Trends in microbiology, 10(3):142–146, 2002.

[9] Paulo Durão and Daniela Güleresi et al. Enhanced survival of ri-fampicin and streptomycin double resistant e. coli inside macrophages. Antimicrobial agents and chemotherapy, pages AAC–00624, 2016.

[10] Chih-Yuan Chiang and Ijeoma Uzoma et al. Mitigating the impact of antibacterial drug resistance through host-directed therapies: Current progress, outlook, and challenges. mBio, 9(1):e01932–17, 2018.

[11] Amber M Smith, Jonathan A McCullers, and Frederick R Adler. Mathematical model of a three-stage innate immune response to a pneumococcal lung infection. Journal of theoretical biology, 276(1):106–116, 2011.

[12] Lucy Ternent, Rosemary J Dyson, Anne-Marie Krachler, and Sara Jabbari. Bacterial fitness shapes the population dynamics of antibiotic-resistant and-susceptible bacteria in a model of combined antibiotic and anti-virulence treatment. Journal of theoretical biology, 372:1–11, 2015.

[13] Bruce R Levin, Fernando Baquero, Peter Pierre Ankomah, and Ingrid C McCall. Phagocytes, antibiotics, and self-limiting bacterial infections. Trends in microbiology, 25(11):878–892, 2017.

[14] Gennady Bocharov and Burkhard Ludewig et al. Underwhelming the immune response: effect of slow virus growth on cd8+-t-lymphocyte responses. Journal of virology, 78(5):2247–2254, 2004.

[15] Dominik Wodarz. Mathematical models of immune effector responses to viral infections: Virus control versus the development of pathology. Journal of computational and applied mathematics, 184(1):301–319, 2005.

[16] Stanca M Ciupe and Ruy M Ribeiro et al. Modeling the mechanisms of acute hepatitis b virus infection. Journal of theoretical biology, 247(1):23–35, 2007.

[17] Satoru Hamada and Masayuki Umemura et al. Il-17a produced by γδ t cells plays a critical role in innate immunity against listeria monocytogenes infection in the liver. The Journal of Immunology, 181(5):3456–3463, 2008.

[18] P Van den Driessche and James Watmough. Further notes on the basic reproduction number. In Mathematical epidemiology, pages 159–178. Springer, 2008.

[19] Rustom Antia, Bruce R Levin, and Robert M May. Within-host population dynamics and the evolution and maintenance of microparasite virulence. The American Naturalist, 144(3):457–472, 1994.

[20] Richard A Proctor and Christof Von Eiff et al. Small colony variants: a pathogenic form of bacteria that facilitates persistent and recurrent infections. Nature Reviews Microbiology, 4(4):295, 2006.

[21] Ricardo S Ramiro, Henrique Costa, and Isabel Gordo. Macrophage adaptation leads to parallel evolution of genetically diverse escherichia coli small-colony variants with increased fitness in vivo and antibiotic collateral sensitivity. Evolutionary applications, 9(8):994–1004, 2016.

[22] Kyle H Rohde and Diogo FT Veiga et al. Linking the transcriptional profiles and the physiological states of mycobacterium tuberculo-analysis during an extended intracellular infection. PLoS pathogens, 8(6):e1002769, 2012.

